# SETD2 is an actin lysine methyltransferase

**DOI:** 10.1101/2020.04.13.034629

**Authors:** Riyad N.H. Seervai, Rahul K. Jangid, Menuka Karki, Durga Nand Tripathi, Sung Yun Jung, Sarah E. Kearns, Kristen J. Verhey, Michael A. Cianfrocco, Bryan A. Millis, Matthew J. Tyska, Frank M. Mason, W. Kimryn Rathmell, In Young Park, Ruhee Dere, Cheryl L. Walker

## Abstract

SET-domain-containing-2 (SETD2) was identified as the methyltransferase responsible for the histone 3 lysine 36 trimethyl (H3K36me3) mark of the histone code. Most recently, SETD2 has been shown to be a dual-function remodeler that regulates genome stability via methylation of dynamic microtubules during mitosis and cytokinesis. Here we show that actin is a *bona fide* target for methylation by SETD2 *in vitro* and in cells. Antibodies against the SETD2 trimethyl lysine epitope recognize methylated actin, with this methyl mark localizing to areas of active actin cytoskeleton reorganization in migrating cells. Disruption of this methylation activity causes defects in actin polymerization and impairs collective cell migration. Together, these data identify SETD2 as a multifunctional cytoskeletal remodeler regulating methylation and polymerization of actin filaments, and provide new avenues for understanding how defects in SETD2 drive disease via aberrant cytoskeletal methylation.

## INTRODUCTION

SET-domain-containing-2 (SETD2), the human homolog of *Drosophila Set2*, is a lysine methyltransferase characterized by its catalytic SET (<u>S</u>u(var)3-9, <u>e</u>nhancer of zeste, trithorax) domain^1^. SETD2 activity as a chromatin remodeler responsible for trimethylation of histone H3 on lysine 36 (H3K36me3) is well-characterized^2,3^. The SETD2 H3K36me3 post-translational modification (PTM) on chromatin participates in alternative splicing, DNA methylation, transcriptional elongation, DNA damage repair, and polycomb silencing during development^4,5^. Loss of SETD2 and the H3K36me3 chromatin mark is embryonic lethal in *Drosophila*^6^ and mice^7^, and *SETD2* defects have been linked to several diseases including cancer^8-10^ and autism spectrum disorder^11-13^.

Recently, an important role for SETD2 outside the nucleus acting on the cytoskeleton to regulate microtubule dynamics via trimethylation of α-tubulin at lysine 40 (α-TubK40me3) was discovered^14^. The catalytic SET domain was shown to methylate α-tubulin *in vitro*, and in dividing cells α-TubK40me3 localized to spindle microtubules and the distal midbody during mitosis and cytokinesis respectively. Loss of SETD2 and the α-TubK40me3 mark led to genomic instability and defects such as multipolar spindle formation, chromosomal bridges at cytokinesis, micronuclei, polyploidy, and polynucleation; a phenotype specifically linked to its activity as a microtubule methyltransferase^14,15^.

Based on these insights that SETD2 is a dual-function chromatocytoskeletal remodeler, we asked whether SETD2 might have other cytoskeletal targets. We found actin is a *bona fide* target for SETD2, and lysine methylation of actin is a new SETD2-dependent modification of the actin cytoskeleton. Antibodies directed against the SETD2 methyl mark localized to areas of active actin remodeling in migrating cells. We also found that disruption of this SETD2 methylation of actin filaments caused defects in actin polymerization and impaired cell migration.

## RESULTS

### SETD2 interacts with actin in cells

SETD2 was present in both the nuclear and cytoplasmic compartments of interphase cells (**Fig. 1A)**, and could be co-immunoprecipitated with endogenous actin in 786-O (clear-cell renal cell carcinoma) cells as well as mCherry-β-actin expressed in HEK293T cells **(Fig. 1B and fig. S1, A and B)**. To test the relationship between SETD2 and potential actin methylation, we asked if a SETD2 methyl epitope could be detected on actin from SETD2-proficient versus SETD2-deficient cells. Antibodies directed against the trimethyl-lysine SETD2 epitope on histones (anti-Me3^K36^) or α-tubulin (anti-Me3^K40^), and a pan-trimethyl lysine epitope (anti-Me3^Pan^) all recognized actin from SETD2-proficient but not SETD2-deficient cells (**Fig. 1C, and fig. S1C**). Re-expression of a functional truncated SETD2 (tSETD2) in SETD2-deficient HKC (human kidney) cells restored actin methylation in conjunction with restoration of histone methylation **(Fig. S1, D and E**). Staining with SETD2 trimethyl-lysine epitope antibodies localized methylation in cells to areas of active cytoskeletal remodeling and high actin turnover, including dorsal ruffles and lamellipodia at the leading edge of cells **(Fig. S1F)**. Consistent with decreased actin methylation seen in the absence of SETD2, signal intensity of the methyl mark in these regions was greatly reduced in SETD2-deficient cells **(Fig. 1D)**. Polyline profiles generated for regions of strong phalloidin staining (polymerized F-actin filaments) showed a correlation between the presence of this methyl mark and polymerized actin, which was significantly reduced in SETD2-deficient cells (**Fig. 1, E and F**). Taken together, these data show actin in cells is methylated in a SETD2-dependent manner, opening up the possibility that actin is a novel target for SETD2 methylation.

**Figure 1.**
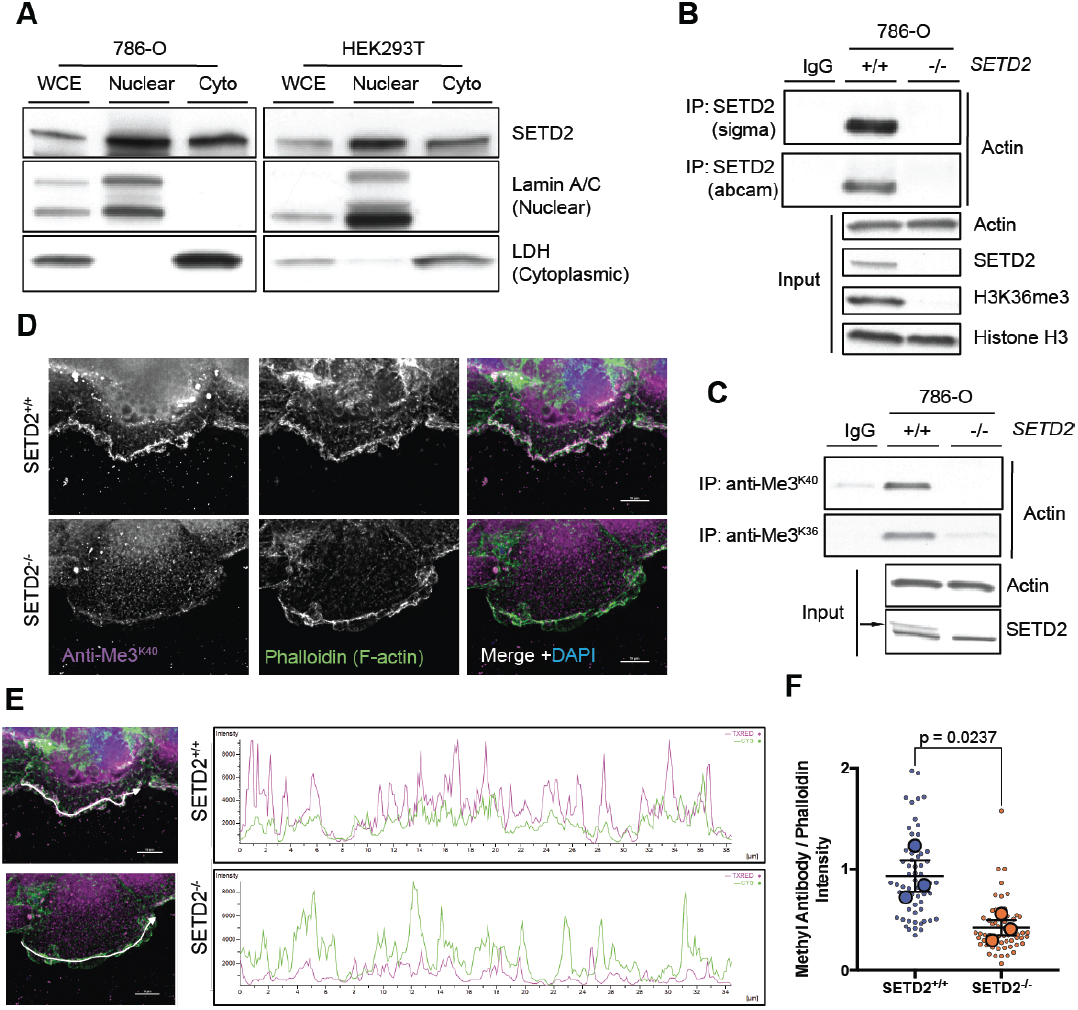
SETD2 binds actin. **(A)** Immunoblot analysis showing localization of SETD2 in whole-cell extracts (WCE) as well as both nuclear and cytoplasmic compartments of 786-O and HEK293T cells. Lamin A/C and lactate dehydrogenase (LDH) used as controls for the nuclear and cytoplasmic (Cyto) fractions, respectively. **(B)** Immunoblot analysis showing co-immunoprecipitation of endogenous SETD2 and endogenous actin in 786-O cells using SETD2 antibodies from two different sources. **(C)** Immunoblot analysis showing actin is methylated in SETD2-proficient but not SETD2-deficient 786-0 cells by immunoprecipitation of endogenous actin using two different antibodies directed against the SETD2 trimethyl-lysine epitope. Data in (A) to (C) are representative of experiments repeated at least three times with similar results. **(D)** Deconvolution microscopy imaging of F-actin using phalloidin (green) and SETD2 methyl epitope-specific antibody (magenta) showing co-localization of the methyl mark with F-actin in 786-O cells. **(E)** Representative intensity profiles of the staining observed with the methyl-specific antibody and phalloidin. Merged images from (D) shown on the left to indicate position for line profiles. **(F)** Quantification of intensity profiles seen in (E). Y-axis represents ratio of Methyl antibody to phalloidin intensity. Each small circle represents a single cell. Large circles represent mean from 20 cells for each independent biological replicate (n=3); *p*-value determined by paired two-tailed t-test.

### SETD2 methylates actin

To directly demonstrate SETD2 could methylate actin, we performed *in vitro* methylation assays based on either incorporation of radiolabeled methyl groups donated by S-adenosylmethionine (^3^H-SAM) or fluorimetric assays based on measurement of SAM consumption. Both assays showed the catalytic SET domain of SETD2 could methylate actins purified from muscle tissues as well as recombinant actin produced in HEK293T cells **(Fig. 2, A and B)**. Recombinant tSETD2 was similarly able to methylate actin purified from cardiac, smooth and skeletal muscle, as well as recombinant actin from HEK293T cells **(Fig. 2 C)**. Trimethyl lysine specific anti-Me3^K36^ and anti-Me3^K40^ antibodies, which immunoprecipitated actin from SETD2-proficient but not SETD2-deficient cells, recognized actin following *in vitro* methylation by SETD2 **(Fig. 2D)**. Interestingly, and consistent with a previous study^16^, SETD2 did not exhibit significant activity against recombinant actin produced in *Escherichia coli* **(Fig. 2, B and C)**, suggesting either proper folding or other modifications may be required to “prime” actin for recognition/methylation. These assays demonstrate SETD2 has intrinsic methylation activity for actin.

**Figure 2.**
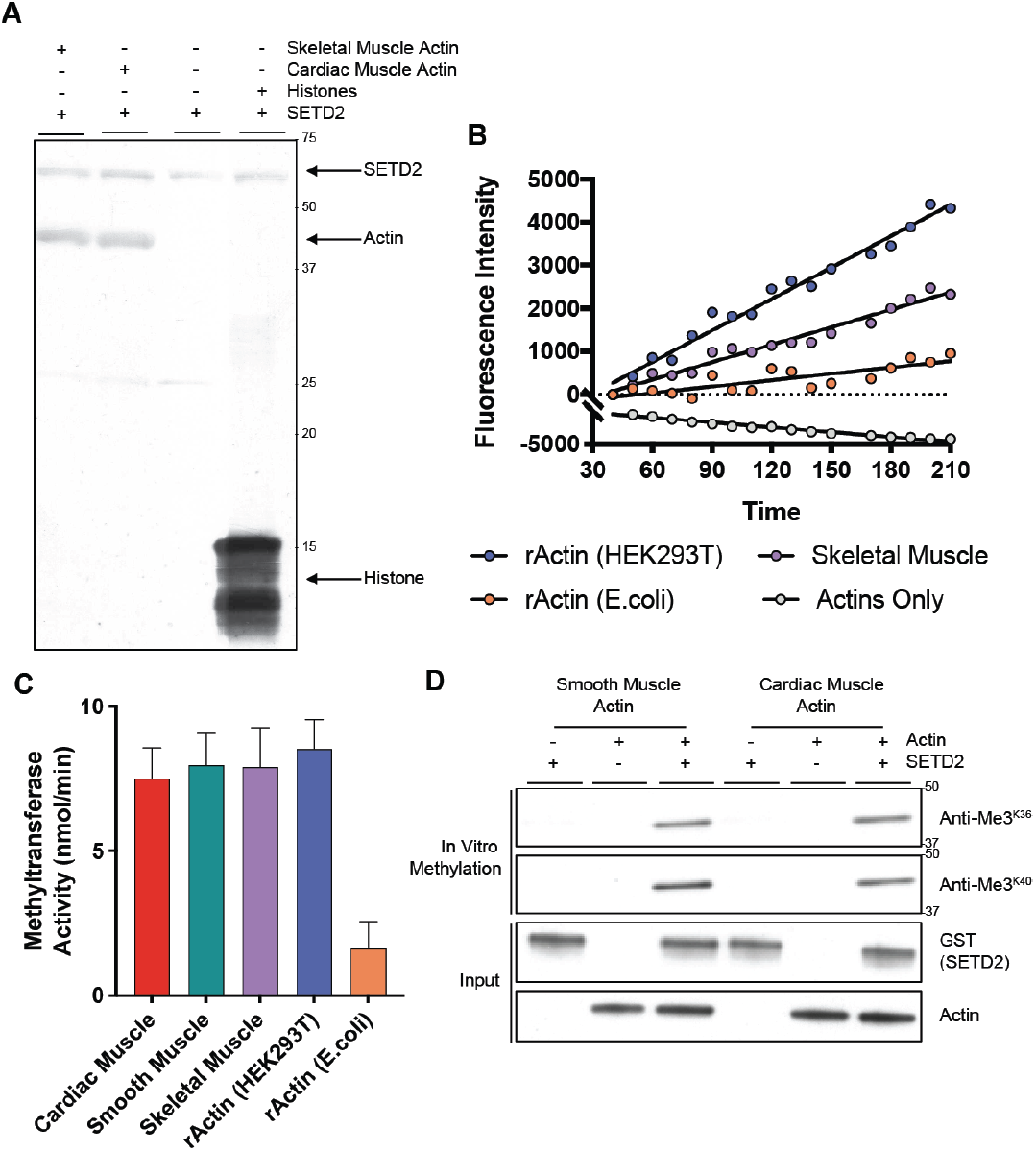
SETD2 methylates actin. **(A)** Autoradiography showing *in vitro* methylation of actin using tritiated S-adenosylmethionine (^3^H-SAM) as methyl group donor and recombinant catalytic SET domain of SETD2. Film shows automethylation of SETD2, as well as histone methylation as positive control. Data are representative of experiments repeated at least three times for similar results. **(B)** Fluorescence-based quantitation of *in vitro* methylation using SETD2 catalytic SET domain (a.a. 1418-1714) with purified skeletal muscle actin (purple), recombinant actin (rActin) from HEK293T cells (blue), rActin from E. coli (orange), or actin proteins alone (grey). **(C)** Fluorescence-based *in vitro* methylation using recombinant tSETD2 (a.a. 1418-2564) with purified cardiac muscle (red), smooth muscle (green), skeletal muscle (purple) actins and rActin from HEK293T cells (blue) or E. coli (orange). Data are mean ± S.E.M (n=2). Y-axis in (B) and (C) plotted after subtracting auto-methylation from samples with SETD2 alone. **(D)** Immunoblot analysis showing recognition of actin proteins by SETD2 methyl-epitope antibodies following *in vitro* methylation with recombinant GST-tagged SETD2 SET domain.

### SETD2 regulates actin polymerization in cells

Based on our findings showing SETD2 is an actin lysine methyltransferase, we sought to understand the impact of SETD2 methylation on the actin cytoskeleton. Using an established biochemical fractionation procedure^17^, we found the ratio of polymerized filamentous F-actin to unpolymerized globular G-actin was significantly reduced in two different SETD2-deficient human cell lines (786-0 and HEK293T), and after acute knockout of *Setd2* in mouse embryo fibroblasts (MEFs), which occurred in the absence of any total difference in overall actin levels (**Fig. 3, A and B**). Re-expression of tSETD2, which rescued methylation, also rescued the polymerization defect in SETD2*-*deficient cells (**Fig. S2, A and B**). These data reveal an actin polymer defect in SETD2-null cells.

**Figure 3.**
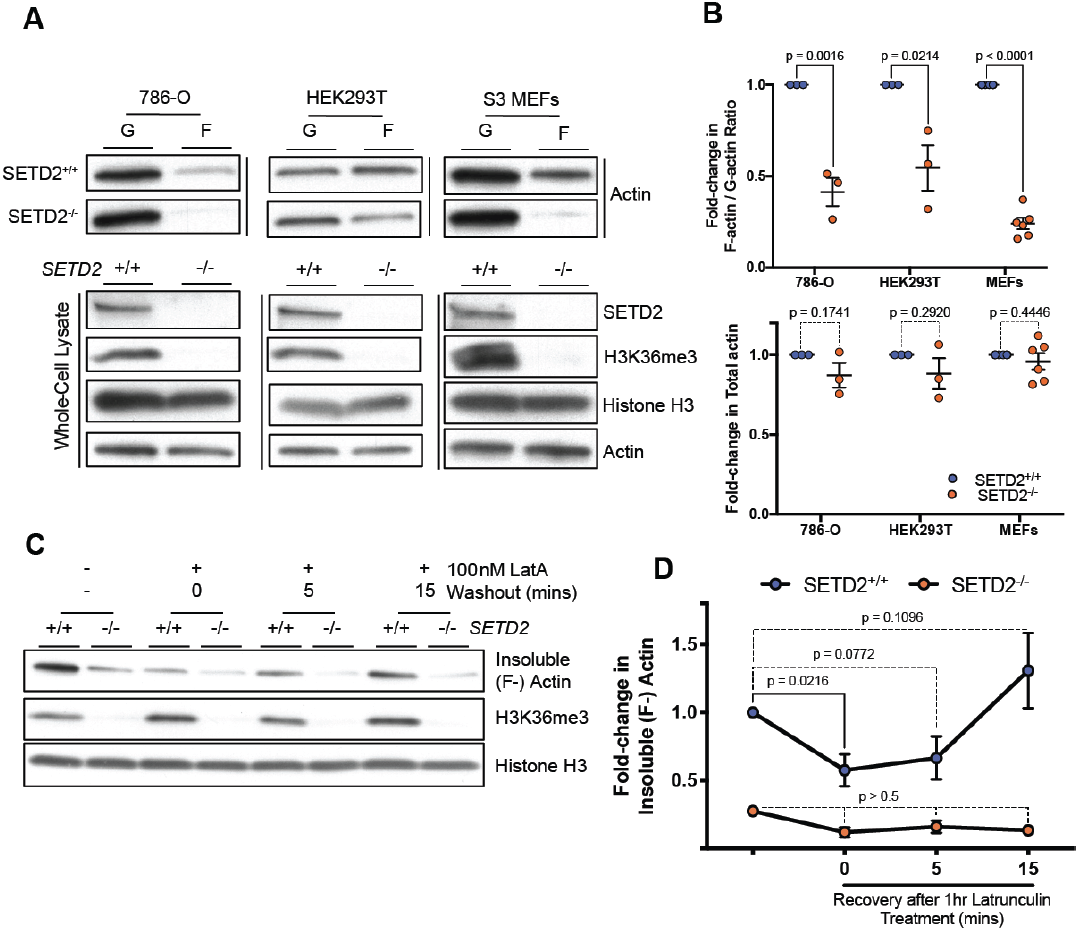
SETD2 regulates actin polymerization in cells. **(A)** Immunoblot analysis showing decreased F-actin in SETD2-deficient 786-O, HEK293T, and mouse embryonic fibroblast (MEF) cells. Whole-cell lysate shows absence of SETD2, associated with the expected loss of histone H3K36me3 methylation. **(B)** Quantitation of F-/G-actin ratio (top) and whole-cell lysate actin (bottom) from data shown in (A). Data are mean ± S.E.M, each circle represents an independent biological replicate (n=3 for 786-O, HEK293T; n=6 for MEF); *p*-value for each cell line determined by unpaired two-tailed t-test. **(C** and **D)** Immunoblot analysis (C) and quantitation (D) showing changes in actin polymerization (F-actin in the insoluble fraction) following washout after treatment with the actin depolymerizing agent Latrunculin A (LatA). Data are mean ± S.E.M. (n=4); *p*-value calculated using ordinary one-way ANOVA with Dunnett’s test for multiple comparisons against DMSO control samples. Absence of H3K36me3 in the insoluble is used as a control to confirm loss of SETD2 in (C).

To investigate the impact of SETD2 loss on actin dynamics, we fractionated cells to isolate the soluble (supernatant) and cytoskeleton-enriched insoluble (pelleted) fractions from cells, and found a reduction in insoluble cytoskeletal F-actin in SETD2-deficient cells (**Fig. S2, C and D**). Treatment with Latrunculin A significantly reduced the insoluble F-actin fraction in SETD2-proficient, but not SETD2-deficient cells in which the polymerized F-actin cytoskeleton was already disrupted. We allowed cells to recover after washout following Latrunculin treatment, and found that polymerized F-actin was restored in SETD2-proficient cells, while cells lacking SETD2 remained deficient for polymerized F-actin in the insoluble fraction (**Fig. 3, C and D**). Interestingly, this defect could be reversed by the polymer-inducing agent Jasplakinolide^18^, which increased F-actin in the insoluble fraction of SETD2-deficient cells to a level comparable to that of SETD2-proficient cells (**Fig. S2, C and D**). We were also able to show the anti-Me3^K40^ SETD2 methyl-lysine epitope and anti-Me3^Pan^ antibodies recognized actin in the insoluble fraction containing polymerized F-actin, and that this immunoreactivity was SETD2-dependent (**Fig. S2, E and F**). Importantly, this fraction lacked any detectable tubulin, confirming that the SETD2 methyl-epitope recognition of actin in this fraction was not due to contamination with the known SETD2 substrate α-tubulin^14^.

Staining with fluorescent phalloidin, which preferentially binds to F-actin polymers^19^, was also used to measure intracellular content of polymerized actin **(Fig. S3A)**. Spectrophotometric analysis as well as flow cytometric analysis of phalloidin-bound actin polymers showed decreased fluorescence intensity (a surrogate for decreased actin polymers) in SETD2-deficient cells **(Fig. S3, B to D)**. High-resolution image stitching of phalloidin-stained SETD2-deficient cells suggested significant compromise of cell-cell contacts, and loss of ubiquitous stress fibers, otherwise present in SETD2-positive cells (**Fig. S4, A and B**). Indeed, super-resolution imaging of the same cells via 3D-structured illumination microscopy (3D-SIM), revealed disorganized actin networks both at the leading edge, as well as lamella, in addition to a decrease in the number of cells displaying robust actin arcs (**Fig. S4C**). Interestingly, SETD2-deficient cells appeared to be associated with a noticeable increase in the appearance of microvilli on the cell surface. **(Fig. S4C)**. Taken together, these biochemical and cellular data point to an actin polymerization defect in cells lacking SETD2.

### SETD2 regulates cell migration

Disruption of actin polymers and localization of the SETD2 methyl mark to the leading edge of cells suggested loss of SETD2 could impact cell migration. Using a scratch wound healing assay, we found SETD2-deficient cells exhibited significantly decreased migration compared to SETD2-proficient cells **(Fig. 4, A and B)**. Treatment with the proliferation inhibitor cytosine arabinoside (AraC), as well as cell counts over 48 hours confirmed decreased cell migration seen in these cells was independent of any effects of SETD2 loss on cell proliferation **(Fig. S5 A and B)**. Expression of tSETD2, which rescued actin methylation and actin polymerization in SETD2-deficient cells, could also rescue the migration phenotype **(Fig. S5, C and D)**. These data link loss of SETD2, actin methylation, and the resulting dysregulation of actin dynamics to defects in cell migration.

**Figure 4.**
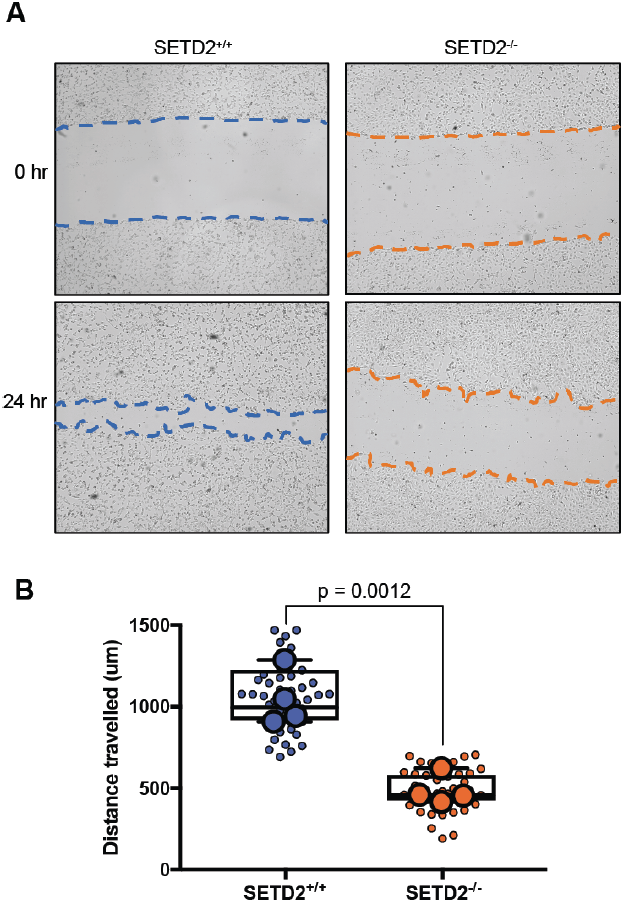
SETD2 regulates cell migration. **(A)** *In vitro* scratch assay illustrating SETD2-proficient migrate faster than SETD2-deficient 786-0 cells at 0 and 24 hours after wound inflection. (**B**) Quantification of scratch assays seen in (A). Small circles each represent an independent measurement across all experiments. Large circles represent mean from a minimum of 9 individual measurements for each independent biological replicate (n=4); *p*-value determined by unpaired two-tailed t-test.

## DISCUSSION

Historically, most studies on SETD2 have focused on the nuclear functions of this enzyme as a histone methyltransferase^4^. The present study demonstrates an extended non-chromatin role for SETD2, and also provides the first example of a lysine methyltransferase capable of modifying and regulating the actin cytoskeleton. Our data show SETD2 methylates actin *in vitro* and in cells, with perturbation of this methylation driving migratory defects seen in cells defective for normal SETD2 function (**Fig. S6**). An earlier study that showed homozygous *Setd2* deficiency was embryonic lethal reported cells from *Setd2*-null mice had disorganized stress fibers and lamellipodia^7^, confirming the link between SETD2 loss and disruption of the actin cytoskeleton.

A recent review of the non-chromatin activity of histone lysine methyltransferases^20^ points out many of these enzymes play roles in the cell that extend beyond their activity on histones. In addition to our data demonstrating SETD2 acts on the actin cytoskeleton, the histone methyltransferase EZH2 has also been found to regulate actin polymerization through interaction with Vav1 and methylation of actin-associated Talin proteins^21,22^. More recently, it was reported that methylation of actin at histidine 73 is mediated by another SET domain-containing protein, the methyltransferase SETD3^16,23^. Whereas histidine methylation of actin has been known to regulate its dynamics for nearly two decades^24^, and Wilkinson *et al*.^16^ found SETD3 to be the only methyltransferase for this site, we show here that lysine methylation is a new regulatory PTM of actin. In that study^16^, *in vitro* methylation experiments found no methyltransferases other than SETD3 (including SETD2) exhibited activity for recombinant actin purified from *E. coli*. However, it is well known this recombinant actin folds poorly^25^, and we also found it was a poor substrate for SETD2 methylation. Furthermore, a new study (in pre-print) shows histidine 73 methylation of actin by the enzyme METTL18 promotes a metastatic response through Src kinase signaling in HER2-negative breast cancer^26^, demonstrating enzymes other than SETD3 can also methylate actin. Regardless, our data, using purified and recombinant actin proteins sourced from models other than *E. coli*, clearly demonstrate that SETD2 methylates actin *in vivo* and *in vitro*, and lysine methylation is a naturally occurring PTM of the actin cytoskeleton.

In conclusion, our findings expand the emerging paradigm that the epigenetic machinery has chromatocytoskeletal activity important in both the nucleus and cytoplasm, coordinately methylating chromatin and the cytoskeleton to regulate the activity of key cellular components during transcription (histones), cell division (microtubules), and cell migration (actin). This new appreciation of SETD2 as a chromatocytoskeletal remodeler will provide new ways to understand how SETD2 defects involved in diseases such as cancer and autism may drive pathogenesis via disruption of cytoskeletal methylation.

## ACKNOWLEDGMENTS

We thank K. Landua and S. Williams (Nikon Instruments Inc.) for assistance with immunofluorescence microscopy experiments. We thank Z. Chriss for assistance with imaging of scratch assay samples. We thank A. Sokac, S.E. McGuire, G. Eisenhoffer, H.C. Hodges, M. Costa-Mattioli, S. Khurana, J. Botas, and P. Msaouel for insightful conversations related to this manuscript. We thank L. Guillen and N. Patel for administrative support.

## AUTHOR CONTRIBUTIONS

C.L.W., R.N.H.S., and I.-Y.P. conceptualized the study. C.L.W., R.N.H.S., I.-Y.P, D.N.T., and R.D. designed experiments, with input from K.J.V., F.M.M., and W.K.R. R.N.H.S. and C.L.W. wrote the manuscript with editorial input from all authors. R.N.H.S. performed experiments and analyzed data contributing to all figures with assistance as noted: R.K.J. and I.-Y.P. assisted with and performed *in vitro* methytransferase assays using recombinant GST-SETD2 (a.a. 1418-1714). M.K. performed and assisted with immunofluorescence assays. D.N.T. assisted with immunoprecipitation, western blotting, siRNA knockdown, and transgene overexpression assays. S.E.K. performed fluorescence-based *in vitro* methyltransferase assays using recombinant tSETD2, with supervision from K.J.V. and M.A.F. B.A.M. performed high-resolution SIM imaging, with supervision from M.J.T. Unless noted, all work was performed under the supervision of C.L.W.

## DATA AND MATERIALS AVAILABILITY

All data are available in the main text and supplementary materials.

## MATERIALS AND METHODS

### Cell Culture

Cell lines were used in the study as previously described^14^. 786-O, HKC, and HEK293T cells were grown in DMEM (Gibco) supplemented with 10% fetal bovine serum (FBS, Sigma, F24429). Cells were incubated at 37 °C and 5% CO_2_ and passaged 2-3 times weekly as required. *Setd2*^flox/flox^ mouse embryonic fibroblast (MEF) cells expressing ER-Cre were cultured in phenol-red-free media (DMEM, high glucose, and HEPES) supplemented with 10% FBS, sodium pyruvate (Gibco) and GlutaMAX (Gibco), with 5 ug/mL blasticidin. MEFs were treated with 2-3 uM 4-hydroxytamoxifen for 3-5 days for efficient knockout of *Setd2*.

siRNA-mediated knockdown in 786-O cells was carried out using 5-10 nM Dharmacon ON-TARGET*plus* siRNAs (GE Healthcare), diluted in siRNA buffer and mixed with DharmaFECT transfection reagent (1:50 in optiMEM reduced serum media). Cells were allowed to incubate with the siRNA mixture for 5-7 hours, and grown for 72-96 hours with passaging as necessary.

Overexpression of mCherry-tagged actin constructs in HEK293T cells was carried out using Lipofectamine 2000 transfection reagent (ThermoFisher) at a ratio of 1:2.5, mixed in optiMEM reduced serum media. Cells were allowed to incubate with the DNA mixture for 5-7 hours, and grown for 48 hours with passaging as necessary.

### Co-immunoprecipitation Assays

Co-immunoprecipitation was performed as previously described^14^, with modifications. In brief, cells were collected at 70-80% confluency and lysed in cold “CST lysis buffer” (20 mM Tris-HCl pH 7.5, 150 mM NaCl, 1 mM EDTA, 1 mM EGTA, 2.5 mM Na_4_P_2_O_7_, 1 mM β-glycerophosphate, 1% Triton X-100) or “IP buffer” (25 mM Tris-HCl, pH 8.0, 300 mM NaCl, 1 mM EDTA, 1% NP-40) supplemented with protease inhibitor cocktail (Roche). Protein concentration was determined using bicinchoninic acid (BCA) protein assay kit as per manufacturer’s instructions (Pierce). Endogenous SETD2 was immunoprecipitated overnight from 500-1000 µg cell extract using anti-SETD2 antibodies (Abcam, or Sigma-Aldrich) and protein A/G agarose beads (Pierce). For the reciprocal IP, endogenous actin was immunoprecipitated using an antibody to actin. In order to identify the interaction between SETD2 and mCherry-tagged, exogenous actin, endogenous SETD2 was immunoprecipitated using whole-cell extracts from HEK293T cells expressing mCherry-ACTB (Addgene plasmid #54966). Similarly, the tagged actin was immunoprecipitated using an mCherry-specific antibody. The appropriate corresponding IgG isotype was used as a negative control.

### Nuclear v. Cytoplasmic Cell Fractionation

Cells were collected in ice-cold PBS, pelleted at 1500 rpm for 5 minutes at 4 °C, and resuspended in an ice-cold hypotonic buffer (10 mM HEPES, pH 7.2, 10 mM KCl, 1.5 mM MgCl_2_, 0.1 mM EGTA, 20 mM NaF, 100 mM Na_3_VO_4_) supplemented with protease inhibitor cocktail. Samples were lysed in a Dounce tissue homogenizer with glass pestle until only nuclei were visible by light microscopy. Samples were centrifuged briefly to separate the supernatant from the nuclei, and the supernatant was centrifuged at 10,000 rpm for 10 minutes and collected as the “cytoplasmic” fraction. The nuclear pellet was lysed in CST lysis buffer, sonicated using a Bioruptor bath sonicator (Diagenode), centrifuged at 10,000 rpm for 10 minutes at 4 °C, and this lysate collected as the “nuclear” fraction. Samples were resolved by SDS-PAGE, and blotted for lamin A/C (CST) or lactate dehydrogenase (Abcam) as markers for the nuclear and cytoplasmic fractions respectively.

### Western Blotting

Samples were prepared at a normalized concentration in phosphate buffer saline (PBS) (3.2 mM Na_2_HPO_4_, 0.5 mM KH_2_PO_4_, 1.3 mM KCl, 135 mM NaCl, pH 7.4) supplemented with protease inhibitor cocktail. All samples were resolved by SDS-PAGE gel electrophoresis using PROTEAN TGX pre-cast gels (Bio Rad) under reducing and denaturing conditions. Resolved proteins were transferred to polyvinyl difluoride (PVDF) membranes at either 300 mAmps for 2-3 hours or at 15 volts overnight. Membranes were blocked with 5% milk in Tris-buffered saline (50 mM Tris-HCl, pH 7.6, 150 mM NaCl) containing 0.5% Tween20 (TBST) for 1 hour at room temperature, and incubated with primary antibody with gentle rocking overnight at 4 °C. Membranes were washed thrice with TBST, incubated with HPR-conjugated secondary antibodies for 1 hour at room temperature, washed thrice with TBST, and developed using Pierce ECL substrate (ThermoFisher) or Amersham ECL Prime (GE Healthcare) for 3 and 5 minutes respectively. Quantitation of immunoblots was performed via densitometric analysis using ImageQuant TL software (GE Healthcare Life Sciences).

### Protein Purification

Glutathione-S-transferase (GST)-tagged protein expressing the catalytic SET domain of SETD2 (a.a. 1418-1714) was purified as previously described^14^. For purification of recombinant tSETD2 (a.a. 1418-2564), tSETD2-Flag plasmid was transfected into HEK293 Freestyle cells with FectoPRO transfection reagent (116-010) and cells were harvested 36-48 hours later at 5,000 rpm for 15 mins (Beckman JLA 8.1, 363563). The pellet was suspended in lysis buffer (50 mM HEPES pH 7.5, 50 mM MgCl_2_, 150 mM NaCl) supplemented with complete protease inhibitor, and cells were lysed with 20 strokes of a dounce homogenizer. This was ultracentrifuged (Beckman Ti70, 337922) at 40,000 rpm for one hour and the supernatant was filtered with 1.0 um glass fiber filter (Pall Laboratory, AP-4527) and incubated with FLAG M2 affinity beads (Sigma Aldrich, A2220) equilibrated in lysis buffer for 3 hours. Beads were rinsed with 3 CV of wash buffer (50 mM NaPi, 150 mM NaCl, 5 mM BME), 3 CV salt buffer (wash buffer at 500 mM NaCl), and again with wash buffer before elution buffer (wash buffer with 300 ng 3x-FLAG peptide (Sigma Aldrich, F4799)) was added and incubated with beads overnight. Eluent was then run over ion exchange column (DEAE Sepharose, GE Life Sciences, 17505501) on a 0-75% salt buffer gradient, and size exclusion chromatography (Superose 6 Increase 10/300, Fisher Scientific, 45-003-210) with gel filtration buffer (50 mM NaPi, 150 mM NaCl, 5 mM BME, 5% glycerol). Fractions were pooled and concentrated down with an Amicon Ultra 100K MWCO (UFC910024) centrifugal filter unit and snap-frozen in liquid nitrogen and stored at −80 °C.

The following commercially available proteins were used in this study: recombinant SETD2 (a.a. 1418-1714) protein (Active Motif, 31358), rabbit skeletal muscle actin (Cytoskeleton, AKL95), bovine cardiac muscle actin (Cytoskeleton, AD99), chicken gizzard smooth muscle actin (Cytoskeleton, AS99), beta actin (NM_001101) human recombinant protein (Origene, TP303643), beta actin (NM_001101) human recombinant protein (Origene, TP720518).

### *In Vitro* Methyltransferase Assays

The intrinsic capacity of SETD2 to methylate actin was demonstrated using *in vitro* methyltransferase reaction assays. To visualize actin methylation via autoradiography, actins were incubated with GST-SETD2 (a.a. 1418-1714) for 3 hours at 37 °C in the presence of tritiated S-adenosylmethionine (^3^H-SAM). Samples were resolved using SDS-PAGE and transferred to PVDF membranes. The signal was amplified using EN3HANCE spray (Perkin Elmer), and detected following incubation with X-ray film for 2-6 weeks. For fluorescence-based assays, the activity of GST-SETD2 (a.a. 1418-1714) and tSETD2-Flag (a.a. 1418-2564) was measured over 3-4 hours using a Methyltransferase Fluorescence Assay Kit (Cayman Chemical, 700150), a continuous enzyme-coupled assay that continuously monitors SAM-dependent methyltransferase activity^27,28^. Readout fluorescence via resorufin was analyzed with an excitation wavelength of 540 nm and an emission wavelength of 590 nm using plate reader. For assays with GST-SETD2, fluorescence was calculated after subtracting auto-methylation signal from samples with SETD2 only. For assays with tSETD2-Flag, a standard curve of resorufin concentration and fluorescence was used to determine time-dependent fluorescence. The initial velocities of these curves were obtained by linear regression to obtain methyltransferase activity in nmol/min using Prism (GraphPad). To detect actin methylation via immunoblotting, 10 uL reactions containing 2 ug actin were incubated with 1 ug GST-SETD2 (a.a. 1418-1714) in an *in vitro* methyltransferase buffer (50 mM Tris-HCl, 2 mM MgCl_2_, 0.02% Triton-X-100, 1 mM DTT, pH 8.6) overnight at 37 °C in the presence of non-tritiated SAM (New England Biolabs). Samples were resolved via SDS-PAGE, and immunoblotted with antibodies overnight at 4 °C followed by secondary antibody incubation for 2 hours at room temperature. Samples with SETD2 alone or actin alone were used as negative controls.

### Immunocytochemistry

Cells were seeded overnight on glass coverslips in a 6-well plate. Samples were fixed in 4% paraformaldehyde in PBS for 30 minutes at 37 °C, permeabilized with 0.5% Triton-X-100 in PBS for 20 minutes, blocked with 3.75% BSA in PBS for 1 hour, and incubated with primary antibody (1:1000) with gentle rocking at 4 °C overnight. Cells were then washed with PBS, incubated with anti-rabbit secondary antibody (1:2000) for 2 hours, and counterstained with DAPI (1:4000) and phalloidin (1:1000) for 10 minutes following post-fixation with 4% paraformaldehyde. Coverslips were rinsed and mounted on glass slides with SlowFade antifade mounting medium (ThermoFisher). All steps were carried at our room temperature unless mentioned otherwise. Immunostained slides were imaged using a CFI Plan Apochromat Lambda 60X oil, 1.4 NA objective and DS-Qi2 camera mounted on a Nikon Eclipse Ti2-E inverted microscope system (Nikon) equipped for standard phase-contrast and epifluorescence, as well as for deconvolution. Image acquisition was carried using an Andor Zyla 4.2+ sCMOS high-sensitivity monochrome camera and was driven by Nikon NIS-Elements Advanced Research (AR) image acquisition and analysis software. Images were processed using advanced deconvolution modules for improved image quality.

For super-resolution imaging of actin, cells were grown on coverslips washed with HCl overnight, and stained with phalloidin (1:40) for 2 hours at 37 °C. Image stitching was accomplished through the use of an inverted epi-fluorescence microscope equipped with a G-2E/C widefield filter cube (λ_ex_:540/25, λ_em_:620/60), automated (and encoded) scanning stage, and 20x Plan Apo Lambda 0.75 NA objective lens (Nikon). The system was additionally outfitted with piezo-based axial positioning (Mad City Labs), LED illumination (Lumencor), and Flash 4.0v3 sCMOS (Hamamatsu Photonics). NIS-Elements was used for both acquisition and analysis (Nikon Instruments, Inc.). For each coordinate (field-of-view) visited, image stacks were acquired to cover the required depth of the cell monolayer. Subsequently, stacks were collapsed through focus-stacking in software, and montage was created. Structured Illumination Microscopy (SIM) was accomplished in 3D-SIM mode on a Nikon Instruments N-SIM, equipped with an Apo TIRF 100x SR 1.49NA objective, 561nm laser, and DU-897 EMCCD camera (Andor). Images presented herein are maximum intensity projections after image stacks were first acquired (5 phase shifts and 3 rotations of diffraction grating, 120nm/axial step) and subsequent stack reconstruction in NIS-Elements software (Nikon Instruments, Inc.). Other than linear intensity scaling, no further image processing was performed post-reconstruction for image panels.

### F-actin to G-actin Ratio

Cell fractionation was performed as previously described^17^, with slight modification, to discriminate between globular (G-) and polymerized filamentous (F-) actin based on the observation that polymerized F-actin is insoluble whereas G-actin is soluble. Cells were collected at 70-80% confluency in ice-cold PBS, pelleted at 500 x g for 5 minutes at 4 °C, and lysed in an ice-cold lysis buffer (10 mM K_2_HPO_4_, 100 mM NaF, 50 mM KCl, 2 mM MgCl_2_, 1 mM EGTA, 0.2 mM DTT, 0.5% Triton-X-100, 1 mM sucrose, pH 7.0). Samples were pipetted repeatedly and vortexed, and centrifuged at 15,000 x g for 10 minutes at room temperature. The supernatant was collected for measurement of G-actin. The insoluble F-actin in the pellet was washed with lysis buffer to remove any residual G-actin, re-suspended in equal volumes of lysis buffer and a second buffer (1.5 mM guanidine hydrochloride,1 mM sodium acetate, 1 mM CaCl_2_, 1 mM ATP, 20 mM Tris-HCl, pH 7.5) and incubated on ice for 20 minutes with gentle mixing every 5 minutes to solubilize polymerized F-actin. The samples were centrifuged at 15,000 x g for 10 minutes at room temperature, and the supernatant was collected for measurement of F-actin. Samples were proportionally loaded and resolved by SDS-PAGE, immunoblotted with an actin antibody.

Alternatively, cells were collected in ice-cold PBS, pelleted at 1500 rpm for 5 minutes at 4 °C, and lysed in a “Solution A” (10 mM Tris-HCl pH 8.0, 3 mM CaCl_2_, 2 mM MgOAc, 0.1 mM EDTA, 320 mM Sucrose, 1 mM DTT, 0.2% NP40) supplemented with protease inhibitor cocktail on ice for 10 minutes. Samples were briefly centrifuged to separate the soluble supernatant and insoluble pellet. The supernatant was centrifuged at 10,000 rpm for 10 minutes at 4 °C. The pellet was washed twice with a “Solution B” (same as Solution A, minus DTT and NP40), resuspended in cold CST lysis buffer, sonicated using a Bioruptor bath sonicator (Diagenode), and centrifuged at 10,000 rpm for 10 minutes at 4 °C. The resulting lysates constitute the soluble G-actin (supernatant) and insoluble F-actin (pellet) fractions of the cell. The insoluble F-actin fraction samples were resolved by SDS-PAGE and blotted with an actin antibody. Samples were normalized to histone H3 as a marker for the insoluble fraction. For immunoprecipitation, 500 ug of normalized lysate was used from the soluble and insoluble fractions for incubation with antibodies as indicated.

### Phalloidin Binding Assay

Fluorimetric analysis of phalloidin binding is based on the observation that phalloidin binds to polymerized (F-) actin only, and thus serves as a readout of intracellular actin polymers, and was performed as previously described^19^. Cells were plated in 24-well dishes and fixed at 70-80% confluency with 4% paraformaldehyde in PBS for 20-30 minutes, and permeabilized in 0.5% Triton X-100 in PBS for 10 minutes. Wells were incubated with Alexa Fluor 568 (AF568) phalloidin (Thermo Fisher) at varying concentrations in PBS for 20 minutes, washed several times with PBS quickly, and the bound phalloidin was extracted from each well using a 0.1 N NaOH solution. The fluorescence intensity for each phalloidin concentration was measured in duplicate using a spectrophotometer with excitation and emission wavelengths of 575 nm and 605 nm respectively. For unstained controls, a 10x excess of unconjugated phalloidin (Sigma) was added in order to obtain a reading for non-specific binding, and this was subtracted from fluorescent readings at each concentration in order to yield fluorescence as a result of specific binding to F-actin. Cells were stained with DAPI following phalloidin extraction and imaged using an Operetta Phenix high-content screening system (Perkin-Elmer). Cells were counted in 3-6 wells for each genotype/treatment, and the fluorimetric intensity was normalized to mean cell number.

For flow cytometric analysis of phalloidin binding, cells were grown to 70-80% confluency, trypsinized and collected in media containing 10% FBS, and pelleted at 500 rpm for 5 minutes at 4 °C. Cells were fixed in 4% paraformaldehyde in PBS with 1 mM EDTA (PBSE) for 20-30 minutes, permeabilized in 0.5% Triton X-100 for 10 minutes, and incubated with AF568 phalloidin at varying concentrations (with or without 10x excess unconjugated phalloidin) for 20 minutes. Cells were washed twice with PBSE between each step. The final samples were re-suspended using PBSE in BD Falcon test tubes with strainer caps, and run on a BD LSRFortessa cell analyzer (BD Biosciences) within 1 hour of preparation. A total of 10,000 events were acquired for each sample, subjected to doublet discrimination, and single-color compensation performed using the 10x unconjugated sample intensity. Samples were analyzed using FlowJo (BD Biosciences).

### *In Vitro* Migration (Scratch) Assay

Quantitative measurement of cell migration was conducted as previously described^29^. In brief, cells were seeded overnight to 90-100% confluence in a 6-well plate. Scratches were made using a 1000 µL pipette tip, with each well washed twice with PBS to remove non-adherent cells. Fresh media was added to the wells, and 9-12 designated points were measured at 0 hrs. The same designated points were measured again after incubation for 24 hrs. Images were analyzed using “Manual Measurement > Length” feature on NIS-Elements software (Nikon). The distance travelled was computed from the difference in scratch width at 0 and 24 hrs.

### *In Vitro* Proliferation Assay

Quantitative measurement of cell proliferation was measured by counting cells over 48 hours (the total duration of the scratch assay migration experiments). 50,000 cells were seeded triplicate in 6-well plates; at 24h and 48h, cells were trypsinized, stained with trypan blue, and manually counted twice each using a hemocytometer.

### Antibodies

The following primary antibodies were used in this study: Actin (C4, mouse, Santa Cruz, sc-47778), Actin (C4, mouse, Millipore, MAB1501), Actin (13E5, rabbit, Cell Signaling, 4970S), GST (mouse, Santa Cruz, sc-138), Histone H3 (D1H2, rabbit, Cell Signaling, 4620S), Histone H3K36me3 (rabbit, Active Motif, 61101), Lamin A/C (rabbit, Cell Signaling, 2032S), LDH (rabbit, Abcam, ab47010), mCherry (16D7, rat, ThermoFisher, M11217), mCherry (rabbit, Abcam, ab167453), mCherry (mouse, Novus, NBP1-96752), Pan anti-trimethyllysine (rabbit, PTM Biolabs, PTM-601), SETD2 (rabbit, Abcam, ab31358), SETD2 (rabbit, Abclonal, A3194), SETD2 (rabbit, Invitrogen, PA5-83615), SETD2 (rabbit, Sigma, HPA-042451), SETD3 (rabbit, Abcam, ab176582), Tubulin (DM1A, mouse, Santa Cruz, sc-32293). Anti-Me3^K40^ methyl-tubulin antibodies were generated as previously described^14,30^.

The following secondary antibodies were used in the study: anti-rabbit HRP (mouse, Santa Cruz, sc-2357), anti-mouse HRP (goat, Santa Cruz, sc-2005), anti-mouse HRP (goat, Bio-Rad, 1706516), anti-rat HRP (goat, Santa Cruz, sc-2065).

The following normal IgG isotype immunoprecipitation controls were used in the study: mouse (Santa Cruz, sc-2025), rat, (Santa Cruz, sc-2026), rabbit (Santa Cruz, sc-2027), rabbit (Abcam, ab37415).

## SUPPLEMENTAL FIGURE LEGENDS

**Figure S1.**
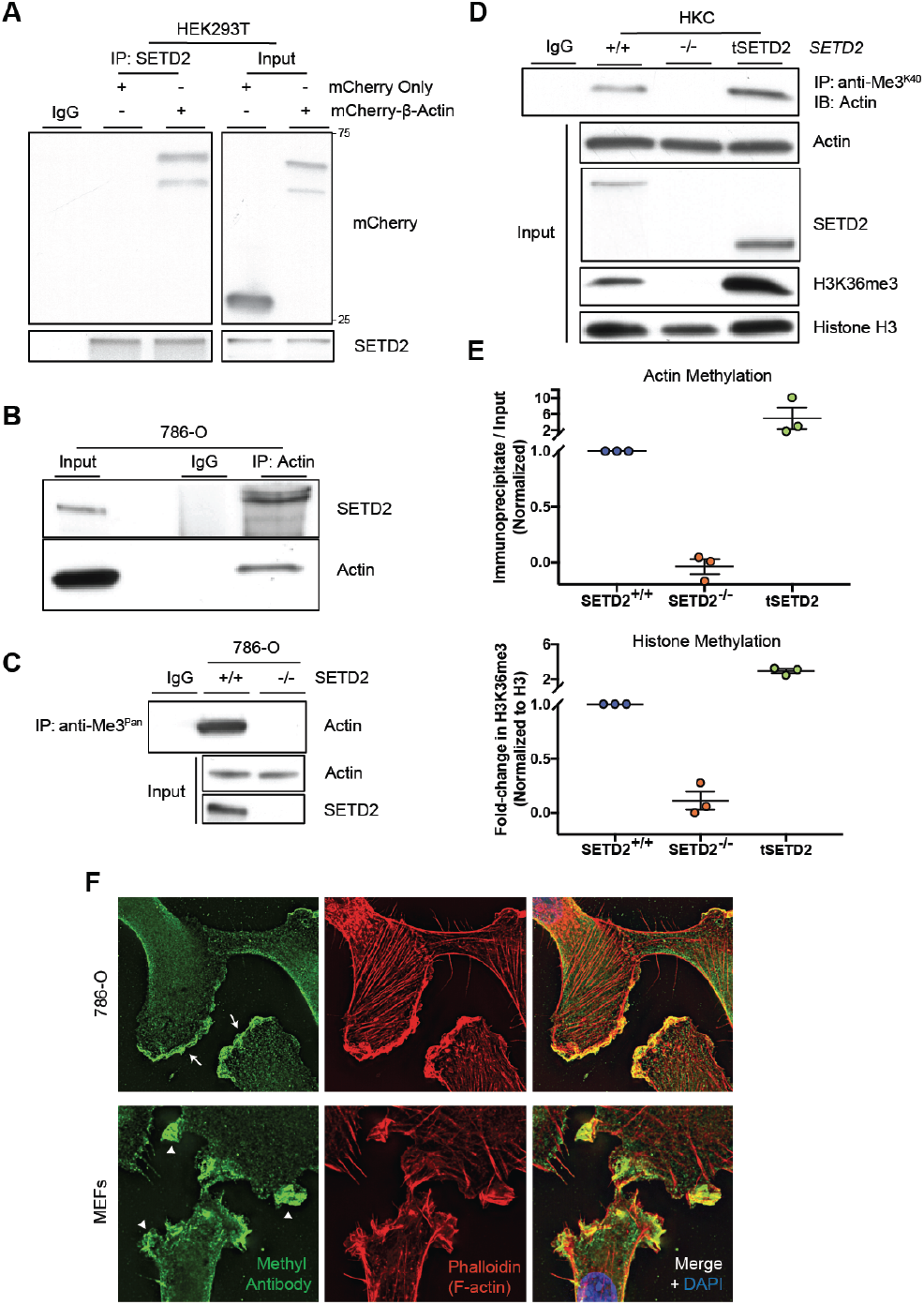
Interaction between SETD2 and actin. **(A)** Immunoblot analysis showing co-immunoprecipitation of endogenous SETD2 with mCherry-β-actin in HEK293T cells. **(B)** Co-immunoprecipitation of endogenous actin with endogenous SETD2 in 786-O cells (reciprocal co-IP from data shown in in fig. 2B). **(C)** Immunoprecipitation of actin from SETD2-proficient but not SETD2-deficient 786-O cells using a pan-trimethyl lysine (anti-Me3^Pan^) antibody. Data in (A) to (C) are representative of experiments repeated at least three times with similar results. **(D** and **E)** Immunoblot (D) and quantitation (E) showing immunoprecipitation of actin from using anti-Me3^K40^ SETD2 methyl-epitope antibody in SETD2-proficient but not SETD2-deficient HKC cells, and rescued by re-expression of tSETD2. Presence or absence of the H3K36me3 histone methyl mark used as a control for SETD2 methyltransferase activity. Data are mean ± S.E.M. for each biological replicate (n=3). **(F)** Deconvolution microscopy imaging of 786-O and mouse embryonic fibroblast (MEF) cells showing localization of the SETD2 methyl epitope (green) to areas of high actin turnover, including ruffles (white arrowheads) and lamellipodia (white arrows) at the leading edge of cells. F-actin stained using phalloidin (shown in red).

**Figure S2.**
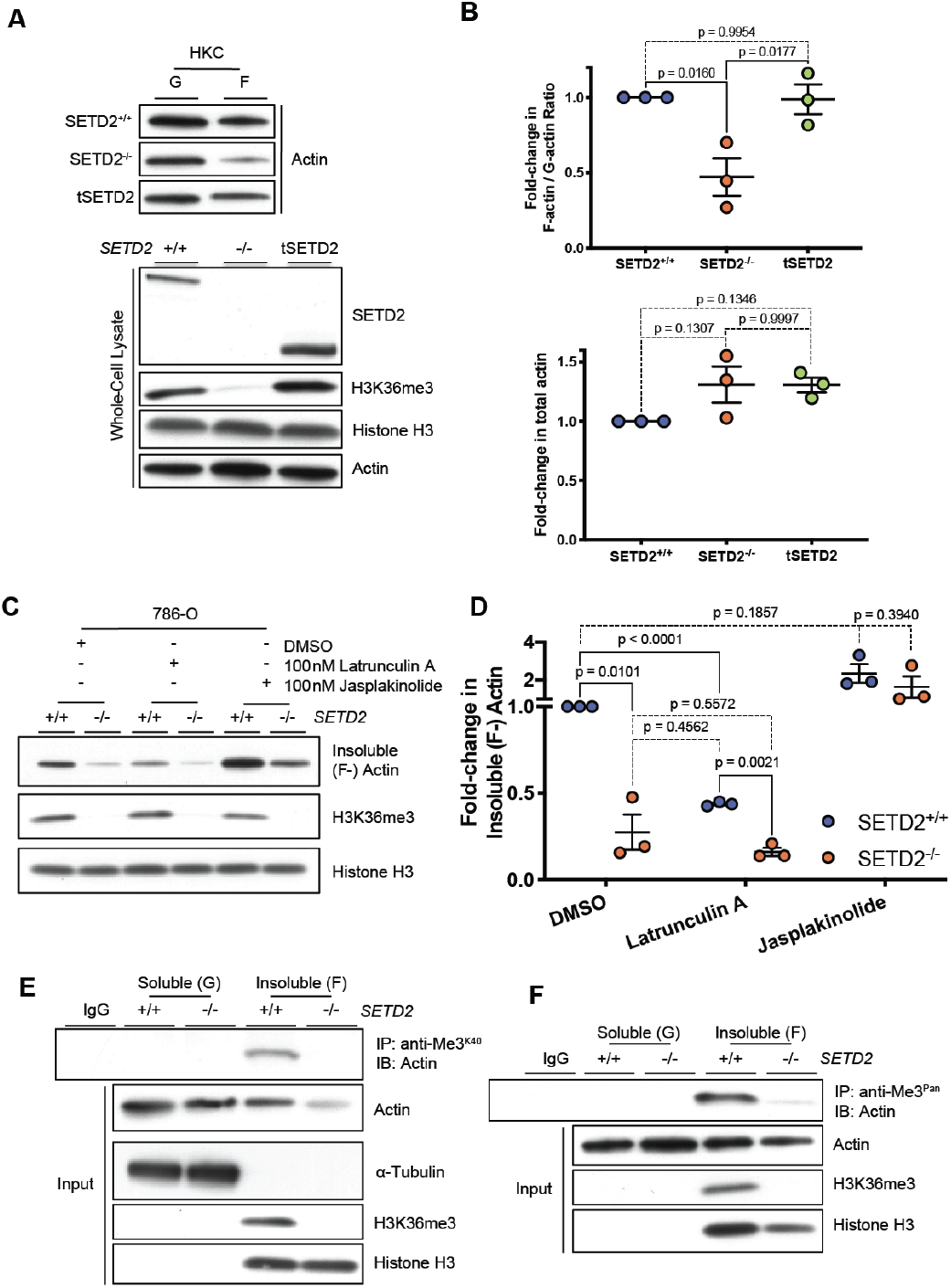
Biochemical evidence for actin defect in SETD2-deficient cells. **(A** and **B)** Immunoblot analysis (A) and quantitation (B) of F-/G-actin ratio in HKC SETD2-proficient, SETD2-deficient, and cells rescued with tSETD2. Data are mean ± S.E.M. (n=3); *p*-value determined by ordinary one-way ANOVA with Tukey’s HSD test for multiple comparisons. **(C)** Immunoblot analysis showing changes in insoluble F-actin in 786-O cells treated with the depolymerizing agent Latrunculin A or the actin polymerizing agent Jasplakinolide. H3K36me3 is used as a control to confirm loss of SETD2. **(D)** Quantitation of changes in insoluble actin fraction shown in (C). Data are mean ± S.E.M. (n=3); *p*-value determined by unpaired two-tailed t-tests with Holm-Šidák correction for multiple comparisons. **(E** and **F)** Loss of the actin methyl mark from polymerized actin in SETD2-deficient cells shown by immunoprecipitation of actin from the insoluble fraction of 786-O cells using SETD2 methyl epitope antibodies anti-Me3^K40^ (E) and anti-Me3^Pan^ (F). Data shown are representative of experiments repeated four times with similar results.

**Figure S3.**
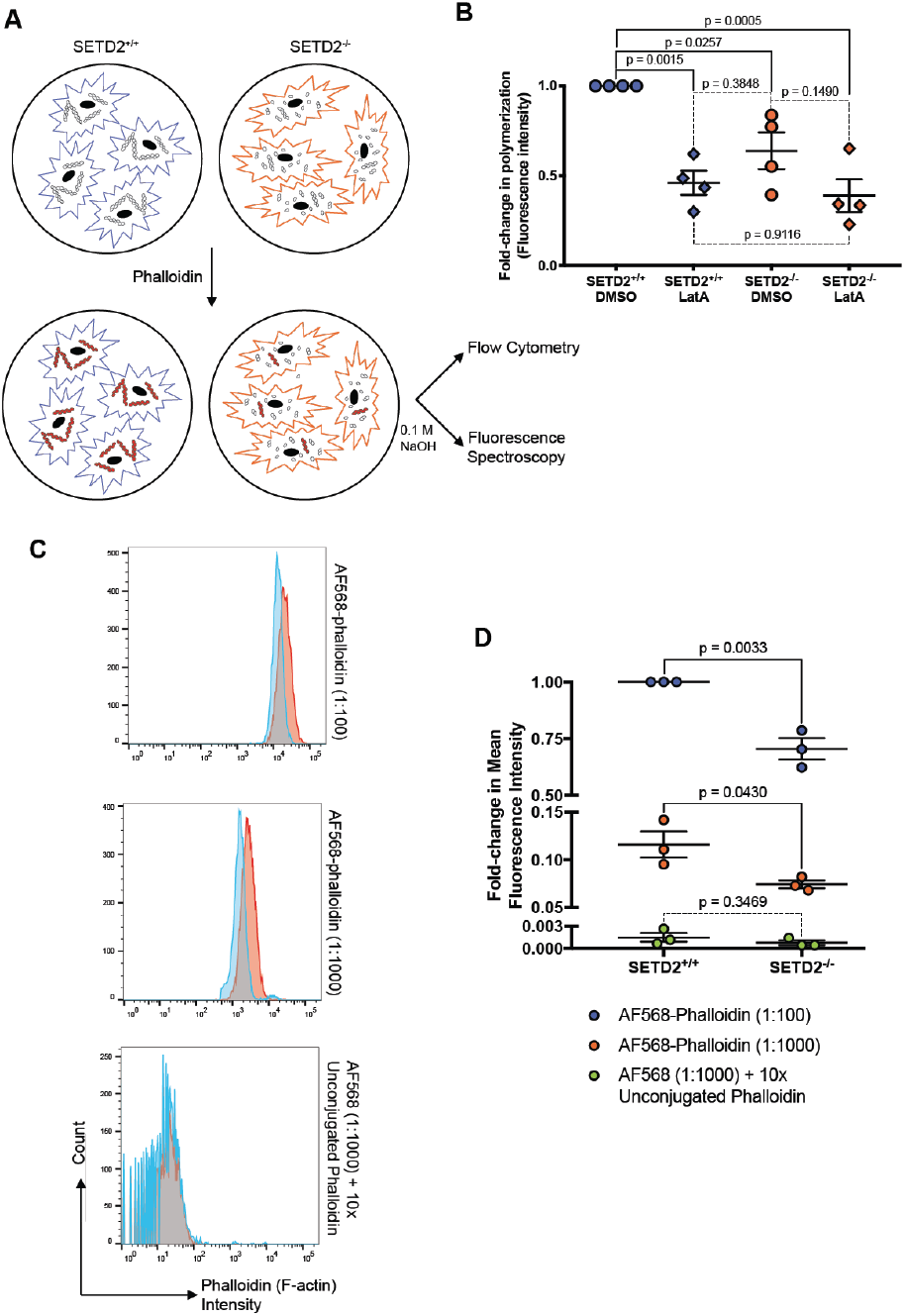
Actin polymerization defect in SETD2-deficient cells. **(A)** Schematic drawing of experiments using fluorescent phalloidin to assess polymerized F-actin content in SETD2-proficient versus-deficient cells. **(B)** Quantitation of spectrophotometric analysis of extracted fluorescent phalloidin from 786-O cells treated with DMSO control or 100 nM Latrunculin A (LatA). Data are mean ± S.E.M. (n=4); *p*-value determined by ordinary one-way ANOVA with Tukey’s HSD test for multiple comparisons. **(C)** Flow cytometry plots showing intensity of fluorescent phalloidin. Data are shown for two concentrations of Alexa Fluor 568 (AF568) conjugated phalloidin (1:100, 1:1000), as well as for samples treated with AF568 phalloidin (1:1000) and 10x unconjugated phalloidin as a control. **(D)** Quantitation of mean fluorescence intensity from plots shown in (C). Y-axis split to reflect magnitude of change in phalloidin intensity between SETD2-proficient and-deficient cells at three different concentrations (1:100, 1:1000, 1:1000 + 10x unconjugated). Data are mean ± S.E.M. (n=3); *p*-value determined by unpaired two-tailed t-test.

**Figure S4.**
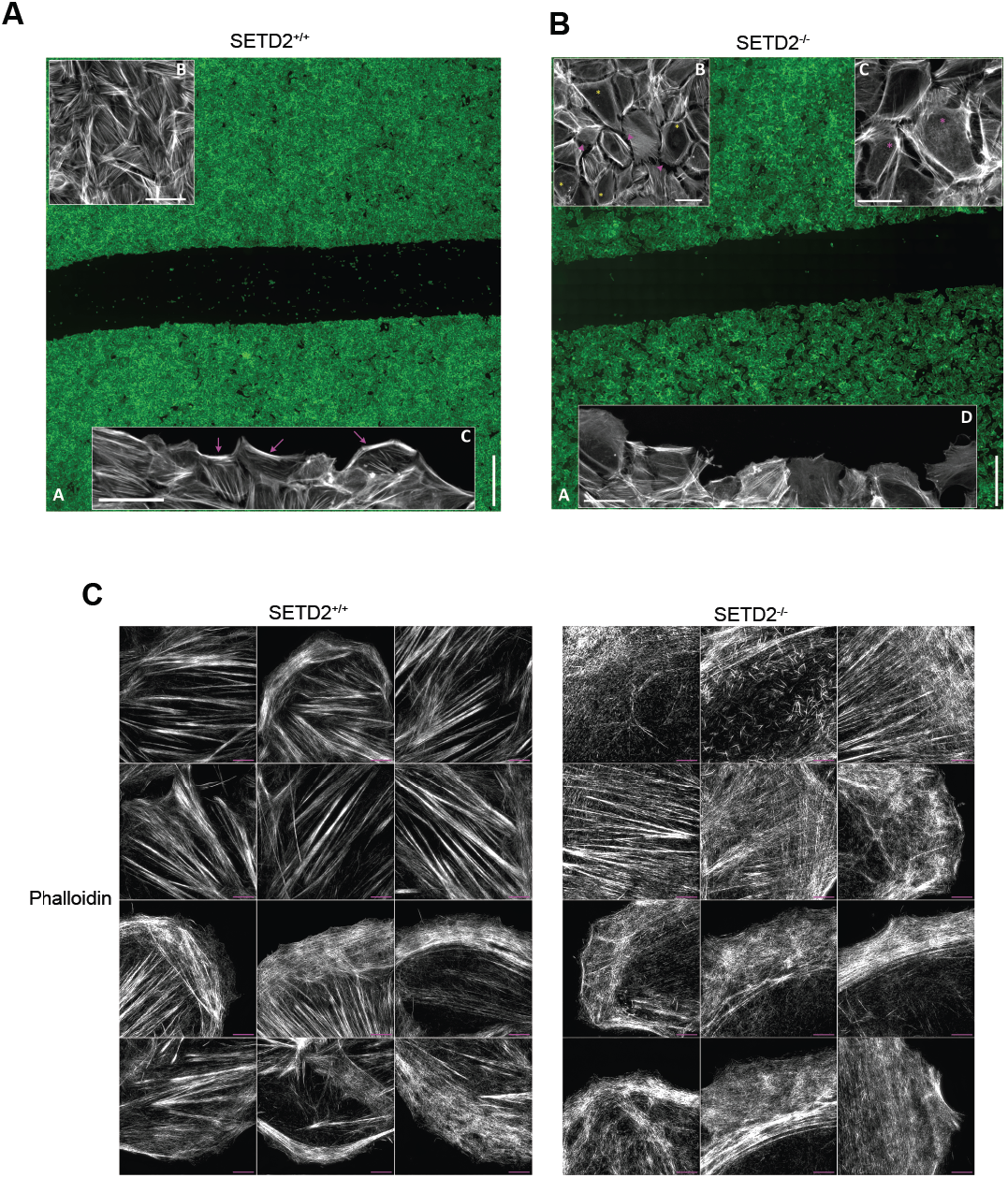
High/Super-Resolution imaging of actin. (**A**) High-Resolution image stitching of SETD2-proficient HKC cells. Sub-panel A shows the ultra-large field-of-view afforded by this automated imaging approach. Sub-panel B displays actual resolution of sub-panel A, to reveal highly confluent, tightly packed monolayer of cells, suggestive of robust cell-cell adhesion. Sub-panel C shows retrograde actin arcs (purple arrows) in leading-edge cells. (**B**) High-resolution imaging of SETD2-deficient HKC cells. Sub-panel A shows ultra-large field-of-view of all cells. Sub-panel B shows dissolution of cell-cell contacts, as seen by increased pericellular space (purple arrowheads), and reduced stress fibers (yellow asterisks) in these cells. Sub-panel C shows apical actin-based protrusions suggestive of microvilli (purple asterisks) not observed in wild-type populations. Sub-panel D shows lack of robust actin arcs seen in wild-type cells, extended lamella, and cells lacking distinct actin signatures at the leading edge. (**C**) Super-Resolution imaging (3D-structured illumination microscopy) revealing robust stress fibers in SETD2-proficient cells (left), relative to the highly disorganized meshwork of actin representative of SETD2-deficient cells, along with appearance of microvilli in cells amongst the population (right).

**Figure S5.**
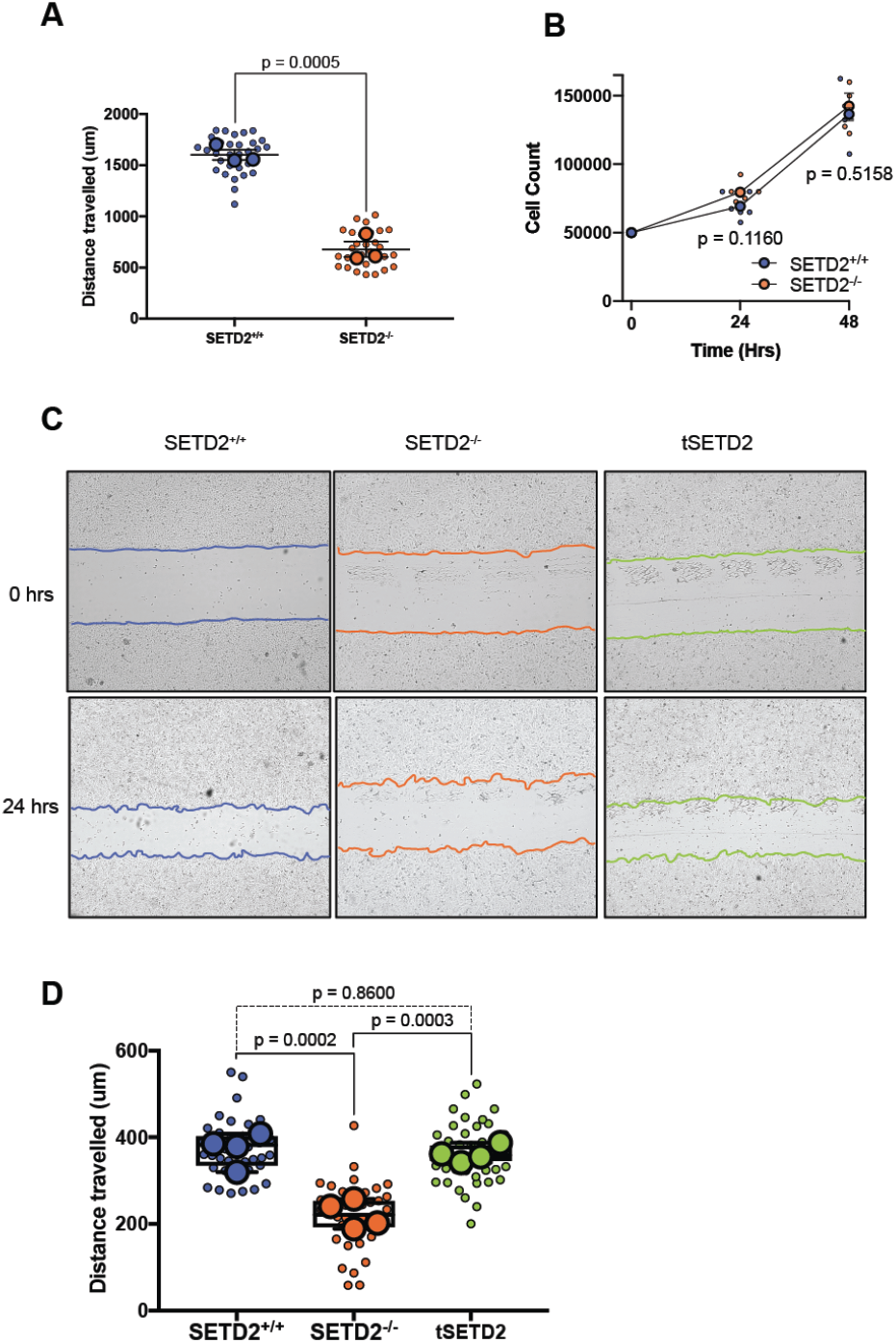
Proliferation-independent migration defect in SETD2-deficient cells. **(A)** Quantitation of scratch assay in 786-O cells treated with cell proliferation inhibitor cytosine arabinoside (AraC). Small circles represent each independent measurement and large circles represent mean from 10 measurements from each independent biological replicate (n=3); *p*-value determined by unpaired two-tailed t-test. **(B)** Cell counts for 786-O cells over 48 hours (duration of the scratch assay from plating to final measurements). Small circles represent each independent measurement and large circles with error bars represent mean ± S.E.M. for each independent biological replicate (n=2); *p*-value determined by unpaired two-tailed t-tests with Holm-Šidák correction for multiple comparisons. **(C)** *In vitro* scratch assay illustrating decreased cell migration in SETD2-deficient versus proficient HKC cells, and rescue of defect by re-expression of tSETD2 at 0 and 24 hours after wound inflection. **(D)** Quantitation of scratch assay seen in (C). Small circles represent each independent measurement and large circles represent mean from a minimum of 9 measurements for each independent biological replicate (n=4); *p*-value determined by ordinary one-way ANOVA with Tukey’s HSD test for multiple comparisons.

**Figure S6.**
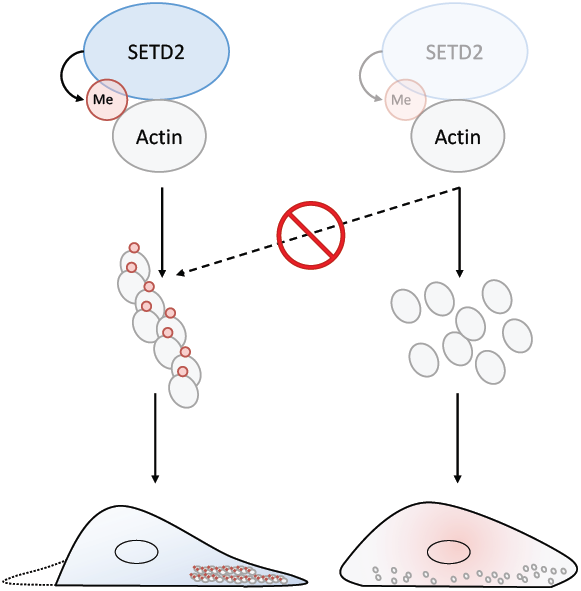
Model. SETD2 methylation is required for normal cellular actin dynamics; loss of SETD2 negatively impacts actin dynamics and cell migration.

## REFERENCES

1 Cheng, X., Collins, R. E. & Zhang, X. Structural and sequence motifs of protein (histone) methylation enzymes. Annu. Rev. Biophys. Biomol. Struct. 34, 267–294, doi:10.1146/annurev.biophys.34.040204.144452 (2005).

2 Edmunds, J. W., Mahadevan, L. C. & Clayton, A. L. Dynamic histone H3 methylation during gene induction: HYPB/Setd2 mediates all H3K36 trimethylation. EMBO J. 27, 406–420, doi:10.1038/sj.emboj.7601967 (2008).

3 Sun, X. J., Wei, J., Wu, X. Y., Hu, M., Wang, L., Wang, H. H., Zhang, Q. H., Chen, S. J., Huang, Q. H. & Chen, Z. Identification and characterization of a novel human histone H3 lysine 36-specific methyltransferase. J. Biol. Chem. 280, 35261–35271, doi:10.1074/jbc.M504012200 (2005).

4 McDaniel, S. L. & Strahl, B. D. Shaping the cellular landscape with Set2/SETD2 methylation. Cell Mol Life Sci, doi:10.1007/s00018-017-2517-x (2017).

5 Wagner, E. J. & Carpenter, P. B. Understanding the language of Lys36 methylation at histone H3. Nat. Rev. Mol. Cell Biol. 13, 115–126, doi:10.1038/nrm3274 (2012).

6 Stabell, M., Larsson, J., Aalen, R. B. & Lambertsson, A. Drosophila dSet2 functions in H3-K36 methylation and is required for development. Biochem. Biophys. Res. Commun. 359, 784–789, doi:10.1016/j.bbrc.2007.05.189 (2007).

7 Hu, M., Sun, X. J., Zhang, Y. L., Kuang, Y., Hu, C. Q., Wu, W. L., Shen, S. H., Du, T. T., Li, H., He, F., Xiao, H. S., Wang, Z. G., Liu, T. X., Lu, H., Huang, Q. H., Chen, S. J. & Chen, Z. Histone H3 lysine 36 methyltransferase Hypb/Setd2 is required for embryonic vascular remodeling. Proc Natl Acad Sci U S A 107, 2956–2961, doi:10.1073/pnas.0915033107 (2010).

8 de Cubas, A. A. & Rathmell, W. K. Epigenetic modifiers: activities in renal cell carcinoma. Nat Rev Urol 15, 599–614, doi:10.1038/s41585-018-0052-7 (2018).

9 Fahey, C. C. & Davis, I. J. SETting the Stage for Cancer Development: SETD2 and the Consequences of Lost Methylation. Cold Spring Harb Perspect Med 7, doi:10.1101/cshperspect.a026468 (2017).

10 Morris, M. R. & Latif, F. The epigenetic landscape of renal cancer. Nat Rev Nephrol 13, 47–60, doi:10.1038/nrneph.2016.168 (2017).

11 Lumish, H. S., Wynn, J., Devinsky, O. & Chung, W. K. Brief Report: SETD2 Mutation in a Child with Autism, Intellectual Disabilities and Epilepsy. J Autism Dev Disord 45, 3764–3770, doi:10.1007/s10803-015-2484-8 (2015).

12 Luscan, A., Laurendeau, I., Malan, V., Francannet, C., Odent, S., Giuliano, F., Lacombe, D., Touraine, R., Vidaud, M., Pasmant, E. & Cormier-Daire, V. Mutations in SETD2 cause a novel overgrowth condition. J Med Genet 51, 512–517, doi:10.1136/jmedgenet-2014-102402 (2014).

13 Marzin, P., Rondeau, S., Aldinger, K. A., Alessandri, J. L., Isidor, B., Heron, D., Keren, B., Dobyns, W. B. & Cormier-Daire, V. SETD2 related overgrowth syndrome: Presentation of four new patients and review of the literature. Am J Med Genet C Semin Med Genet 181, 509–518, doi:10.1002/ajmg.c.31746 (2019).

14 Park, I. Y., Powell, R. T., Tripathi, D. N., Dere, R., Ho, T. H., Blasius, T. L., Chiang, Y. C., Davis, I. J., Fahey, C. C., Hacker, K. E., Verhey, K. J., Bedford, M. T., Jonasch, E., Rathmell, W. K. & Walker, C. L. Dual Chromatin and Cytoskeletal Remodeling by SETD2. Cell 166, 950–962, doi:10.1016/j.cell.2016.07.005 (2016).

15 Chiang, Y. C., Park, I. Y., Terzo, E. A., Tripathi, D. N., Mason, F. M., Fahey, C. C., Karki, M., Shuster, C. B., Sohn, B. H., Chowdhury, P., Powell, R. T., Ohi, R., Tsai, Y. S., de Cubas, A. A., Khan, A., Davis, I. J., Strahl, B. D., Parker, J. S., Dere, R., Walker, C. L. & Rathmell, W. K. SETD2 Haploinsufficiency for Microtubule Methylation Is an Early Driver of Genomic Instability in Renal Cell Carcinoma. Cancer Res 78, 3135–3146, doi:10.1158/0008-5472.CAN-17-3460 (2018).

16 Wilkinson, A. W., Diep, J., Dai, S., Liu, S., Ooi, Y. S., Song, D., Li, T. M., Horton, J. R., Zhang, X., Liu, C., Trivedi, D. V., Ruppel, K. M., Vilches-Moure, J. G., Casey, K. M., Mak, J., Cowan, T., Elias, J. E., Nagamine, C. M., Spudich, J. A., Cheng, X., Carette, J. E. & Gozani, O. SETD3 is an actin histidine methyltransferase that prevents primary dystocia. Nature 565, 372–376, doi:10.1038/s41586-018-0821-8 (2019).

17 Gu, Y. Y., Zhang, H. Y., Zhang, H. J., Li, S. Y., Ni, J. H. & Jia, H. T. 8-Chloroadenosine inhibits growth at least partly by interfering with actin polymerization in cultured human lung cancer cells. Biochemical pharmacology 72, 541–550, doi:10.1016/j.bcp.2006.05.026 (2006).

18 Holzinger, A. Jasplakinolide: an actin-specific reagent that promotes actin polymerization. Methods Mol Biol 586, 71–87, doi:10.1007/978-1-60761-376-3_4 (2009).

19 Cooper, J. A. in The Cytoskeleton: A Practical Approach (ed K. L. & Carraway Carrawayy, C. A. C.) 47–71 (Oxford University Press, 1992).

20 Cornett, E. M., Ferry, L., Defossez, P. A. & Rothbart, S. B. Lysine Methylation Regulators Moonlighting outside the Epigenome. Mol Cell 75, 1092–1101, doi:10.1016/j.molcel.2019.08.026 (2019).

21 Gunawan, M., Venkatesan, N., Loh, J. T., Wong, J. F., Berger, H., Neo, W. H., Li, L. Y., La Win, M. K., Yau, Y. H., Guo, T., See, P. C., Yamazaki, S., Chin, K. C., Gingras, A. R., Shochat, S. G., Ng, L. G., Sze, S. K., Ginhoux, F. & Su, I. H. The methyltransferase Ezh2 controls cell adhesion and migration through direct methylation of the extranuclear regulatory protein talin. Nat Immunol 16, 505–516, doi:10.1038/ni.3125 (2015).

22 Su, I. H., Dobenecker, M. W., Dickinson, E., Oser, M., Basavaraj, A., Marqueron, R., Viale, A., Reinberg, D., Wulfing, C. & Tarakhovsky, A. Polycomb group protein ezh2 controls actin polymerization and cell signaling. Cell 121, 425–436, doi:10.1016/j.cell.2005.02.029 (2005).

23 Kwiatkowski, S., Seliga, A. K., Vertommen, D., Terreri, M., Ishikawa, T., Grabowska, I., Tiebe, M., Teleman, A. A., Jagielski, A. K., Veiga-da-Cunha, M. & Drozak, J. SETD3 protein is the actin-specific histidine N-methyltransferase. Elife 7, doi:10.7554/eLife.37921 (2018).

24 Nyman, T., Schuler, H., Korenbaum, E., Schutt, C. E., Karlsson, R. & Lindberg, U. The role of MeH73 in actin polymerization and ATP hydrolysis. J Mol Biol 317, 577–589, doi:10.1006/jmbi.2002.5436 (2002).

25 Karlsson, R. Expression of chicken beta-actin in Saccharomyces cerevisiae. Gene 68, 249–257 (1988).

26 Kim, H. G., Kim, J. H., Yang, W. S., Park, J. G., Lee, Y. G., Hong, Y. H., Kim, E., Jo, M., Lee, C. Y., Kim, S. H., Sung, N. Y., Yi, Y.-S., Ratan, Z. A., Kim, S., Yoo, B. C., Kang, S.-U., Kim, Y. B., Kim, S., Paik, H.-J., Lee, J. E., Nam, S. J., Parameswaran, N., Han, J.-W. & Cho, J. Y. Metastatic function of METTL18 in breast cancer via actin methylation and Src. bioRxiv, doi:10.1101/831701 (2019).

27 Burgos, E. S., Walters, R. O., Huffman, D. M. & Shechter, D. A simplified characterization of S-adenosyl-l-methionine-consuming enzymes with 1-Step EZ-MTase: a universal and straightforward coupled-assay for in vitro and in vivo setting. Chem Sci 8, 6601–6612, doi:10.1039/c7sc02830j (2017).

28 Dorgan, K. M., Wooderchak, W. L., Wynn, D. P., Karschner, E. L., Alfaro, J. F., Cui, Y., Zhou, Z. S. & Hevel, J. M. An enzyme-coupled continuous spectrophotometric assay for S-adenosylmethionine-dependent methyltransferases. Anal Biochem 350, 249–255, doi:10.1016/j.ab.2006.01.004 (2006).

29 Chowdhury, B., Porter, E. G., Stewart, J. C., Ferreira, C. R., Schipma, M. J. & Dykhuizen, E. C. PBRM1 Regulates the Expression of Genes Involved in Metabolism and Cell Adhesion in Renal Clear Cell Carcinoma. PLoS One 11, e0153718, doi:10.1371/journal.pone.0153718 (2016).

30 Park, I. Y., Chowdhury, P., Tripathi, D. N., Powell, R. T., Dere, R., Terzo, E. A., Rathmell, W. K. & Walker, C. L. Methylated alpha-tubulin antibodies recognize a new microtubule modification on mitotic microtubules. MAbs 8, 1590–1597, doi:10.1080/19420862.2016.1228505 (2016).

